# Widespread Sexual Dimorphism in the Transcriptome of Human Airway Epithelium in Response to Smoking

**DOI:** 10.1101/602854

**Authors:** Chen Xi Yang, Henry Shi, Irving Ding, Cheng Wei Tony Yang, Edward Kyoo-Hoon Kim, Tillie-Louise Hackett, Janice Leung, Don D. Sin, Ma’en Obeidat

## Abstract

Epidemiological studies have shown that female smokers are at higher risk of chronic obstructive pulmonary disease (COPD). Female patients have worse symptoms and health status and increased risk of exacerbations. We determined the differences in the transcriptome of the airway epithelium between males and females at baseline and in response to smoking. We processed public gene expression data of human airway epithelium into a discovery cohort of 211 subjects (never smokers n=68; current smokers n=143) and two replication cohorts of 104 subjects (21 never, 52 current, and 31 former smokers) and 238 subjects (99 current and 139 former smokers. We analyzed gene differential expression with smoking status, sex, and smoking-by-sex interaction and used network approaches for modules’ level analyses. We identified and replicated two differentially expressed modules between the sexes in response to smoking with genes located throughout the autosomes and not restricted to sex chromosomes. The two modules were enriched in autophagy (up-regulated in female smokers) and response to virus and type 1 interferon signaling pathways which were down-regulated in female smokers compared to males. The results offer insights into the molecular mechanisms of the sexually dimorphic COPD risk and presentation potentially enabling a precision medicine approach to COPD.

## INTRODUCTION

Chronic obstructive pulmonary disease (COPD) affects more than 300 million people and is the 3^rd^ leading cause of death worldwide ^1^. In Canada, COPD is the leading cause of hospitalization, accounting for 80,000 hospital admissions per year^2^. While historically men used to smoke more, there has been a marked rise in the prevalence of smoking among women since the 1960’s, leading to a sharp increase in the burden of COPD among women. In the year 2000 the number of women dying of COPD surpassed that of men in the United States ^3^, and in Canada recent estimates suggest that that there are approximately 85% more female COPD patients than male patients ^4^. Our group has reported previously that female smokers experience a heightened risk of COPD, particularly following menopause ^5^. These data have been validated in a study involving 149,075 women and 100,252 men in the UK Biobank ^6^. For the same severity of airflow obstruction, female patients have increased shortness of breath, poorer quality of life and health status and greater functional impairment than male patients^7^. Furthermore, women with COPD are more likely to experience increased risk of exacerbations ^8^.

There is a growing body of evidence from histological, imaging and animal models to support the notion that females have more airways disease and less emphysema compared to male COPD patients^9,10^. In a murine model where male and female (ovariectomized and ovary-intact) mice were exposed to smoke for 6 months, the female mice with intact ovaries had significantly thicker small airway walls but had similar degrees of emphysema compared with male or ovariectomized female mice ^11^.

Females are also thought to have augmented xenobiotic metabolism of inhaled chemicals such as nicotine ^12^. Since some metabolites of nicotine may be toxic, the sex-related differences in xenometabolism maybe an important driver of increased female susceptibility to COPD ^13,14^.

We have shown *in vitro* that females have enhanced ability to produce oxidants in the airways in response to cigarette smoking, leading to increased airway damage ^15,16^.

To further understand the sexual dimorphism in disease risk and response to smoking, we determined whether there were major differences in the transcriptome of the airway epithelium between males and females at baseline and in response to smoking.

## RESULTS

The demographics of the discovery and replication cohorts are described in **Table 1**. The discovery set included 68 never smokers and 143 current smokers while the replication set included 21 never smokers, 31 former smokers and 52 current smokers. There were no statistically significant differences in age or pack-years between female never smokers, female current smokers, male never smokers and male current smokers in the discovery or the replication sets (all Bonferroni adjusted p-values > 0.05). We also used a third dataset (GSE37147) for potential replication although it only contained data on former and current smokers (i.e. there were no never-smokers). However, compared to the discovery or the replication sets, the female current smokers, the subjects in GSE37147 were significantly older (all Bonferroni adjusted p-values < 0.05). The male current smokers in GSE37147 also had increased pack-years compared to those in the discovery or thereplication sets (Bonferroni adjusted p-values < 0.05). The female current smokers in GSE37147 had increased pack-years compared to those in the discovery set (Bonferroni adjusted p-value = 9.04E-03) but not to those in the replication set (Bonferroni adjusted p-value = 0.95).

**Table 1.** Demographics of subjects in the discovery and the replication set. Descriptive statistics are shown as (median, interquartile range). NS = Never Smoker, FS = Former Smoker, CS = Current Smoker.

|  | Discovery Set |  |  |  | Replication Set |  |  |  |  |  | GSE37147 |  |  |  |
| --- | --- | --- | --- | --- | --- | --- | --- | --- | --- | --- | --- | --- | --- | --- |
| Sex | Female<br>n=70, 33.2% |  | Male<br>n=141, 66.8% |  | Female<br>n=25, 24.0% |  |  | Male<br>n=79, 76.0% |  |  | Female<br>n=103, 43.3% |  | Male<br>n=135, 56.7% |  |
| Smoking Status | NS<br>n=26,<br>37.1% | CS<br>n=44,<br>62.9% | NS<br>n=42,<br>29.8% | CS<br>n=99,<br>70.2% | NS<br>n=5,<br>20.0% | FS<br>n=10,<br>40.0% | CS<br>n=10,<br>40.0% | NS<br>n=16,<br>20.3% | FS<br>n=21,<br>26.6% | CS<br>n=42,<br>53.1% | FS<br>n=54,<br>52.4% | CS<br>n=49,<br>47.6% | FS<br>n=85,<br>63.0% | CS<br>n=50,<br>37.0% |
| Age | (38.0,<br>23.5) | (46.0,<br>12.5) | (39.5,<br>11.5) | (45.0,<br>7.5) | (28.0,<br>7.0) | (44.0,<br>11.8) | (40.5,<br>13.5) | (29.5,<br>14.5) | (63.0,<br>18.0) | (47.5,<br>25.3) | (66.0,<br>6.98) | (63.7,<br>6.5) | (66.3,<br>8.7) | (61.1,<br>7.37) |
| Pack years | (0.0,<br>0.0) | (28.5,<br>16.8) | (0.0,<br>0.0) | (26.0,<br>21.8) | (0.0,<br>0.0) | (6.0,<br>25.8) | (24.0,<br>17.1) | (0.0,<br>0.0) | (35.0,<br>52.0) | (25.0,<br>39.4) | (42.0,<br>15.0) | (47.1,<br>15.1) | (47.6,<br>19.9) | (43.2,<br>13.8) |
| COPD status | Yes<br>(n=0) | Yes<br>(n=11) | Yes<br>(n=0) | Yes<br>(n=31) | NA | NA | NA | NA | NA | NA | Yes<br>(n=21) | Yes<br>(n=14) | Yes<br>(n=36) | Yes<br>(n=16) |

### Prediction of sex in the replication set

The best-performing elastic net-regularized logistic regression model consisted of three probes and together they demonstrated 100% cross-validated AUC in the training dataset. These three probes corresponded to *RPS4Y1, DDX3Y* and *KDM5D* genes, which are located on the Y-chromosome. The model was then evaluated in the testing dataset and also showed perfect ability to discriminate with an AUC of 100% (**Supplementary Figure 2A and 2B**). After confirming the model performance, we applied it to the replication set to classify the subjects into 79 males and 25 females (**Supplementary Figure 2C**).

### Differential expression with sex and smoking

As shown in the PCA plot in **Supplementary Figure 1E**, there was clear clustering in the gene expression profiles of the discovery set, the replication set and GSE37147. The differences may be related to different microarray platforms that were used and different generations of the airway that were sampled across the 3 studies

In the discovery set, 8,000 probes, corresponding to 5,466 genes, were differentially expressed between never and current smokers after adjustment for sex, age, COPD, ethnicity and pack-years of smoking (FDR < 0.1) and 735 of these genes were replicated in the replication set (at a threshold of P < 0.05) (**Figure 2A and 2B**). A gene was considered “replicated” if any of the probes showed differential expression between never and current smokers in both the discovery and replication sets with the same direction of effect. The top three replicated genes associated with smoking were *ADH7* (FDR=1.15E-32), *GPX2* (FDR=1.43E-31) and *ALDH3A1* (FDR=8.56E-31), which were all up-regulated in current smokers. We also validated the discovery in a second replication set GSE37147; out of the 735 replicated genes, 526 were further replicated in GSE37147 in former versus current smokers.

**Figure 1.**
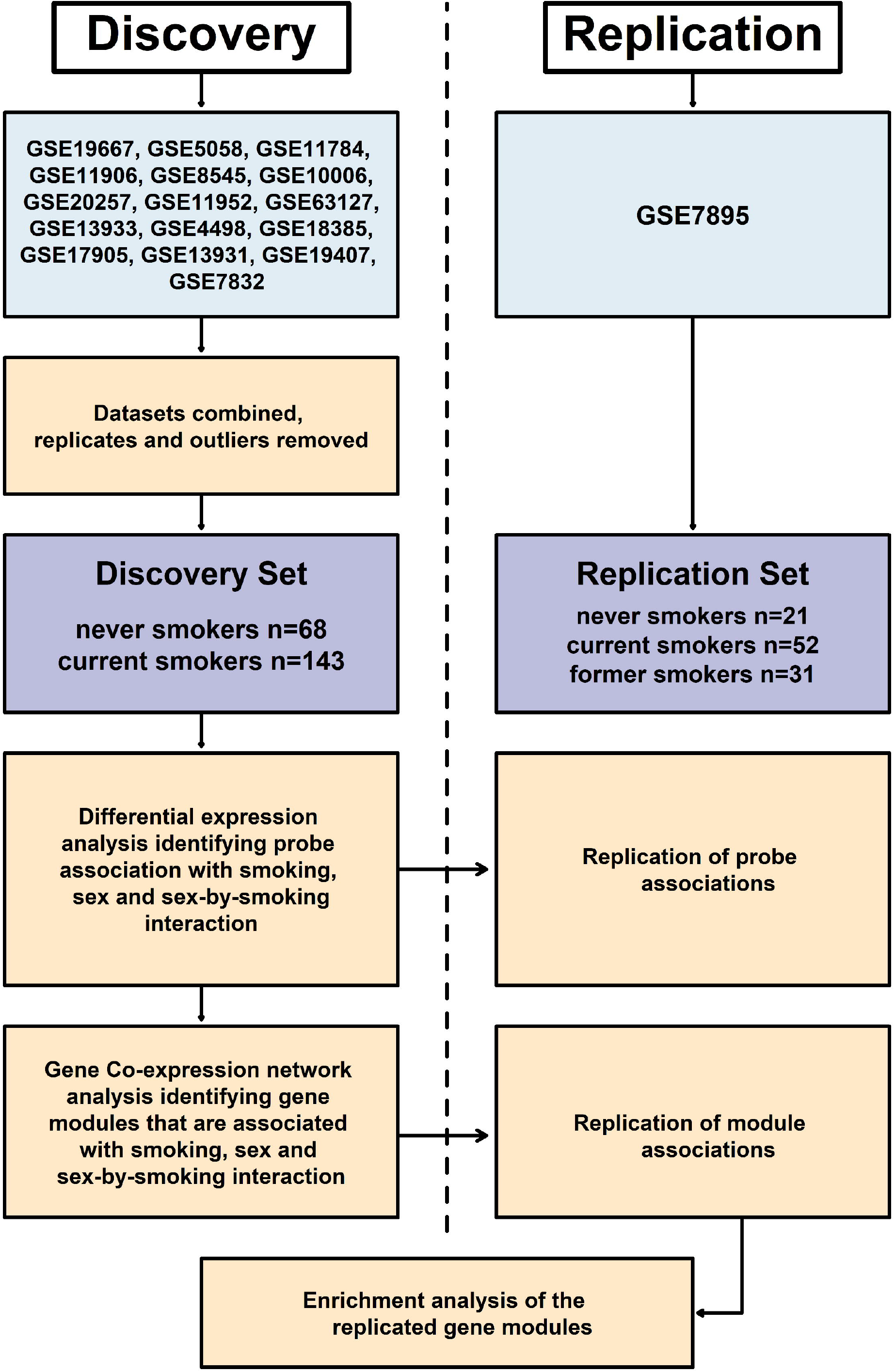
Overall study design

**Figure 2.**
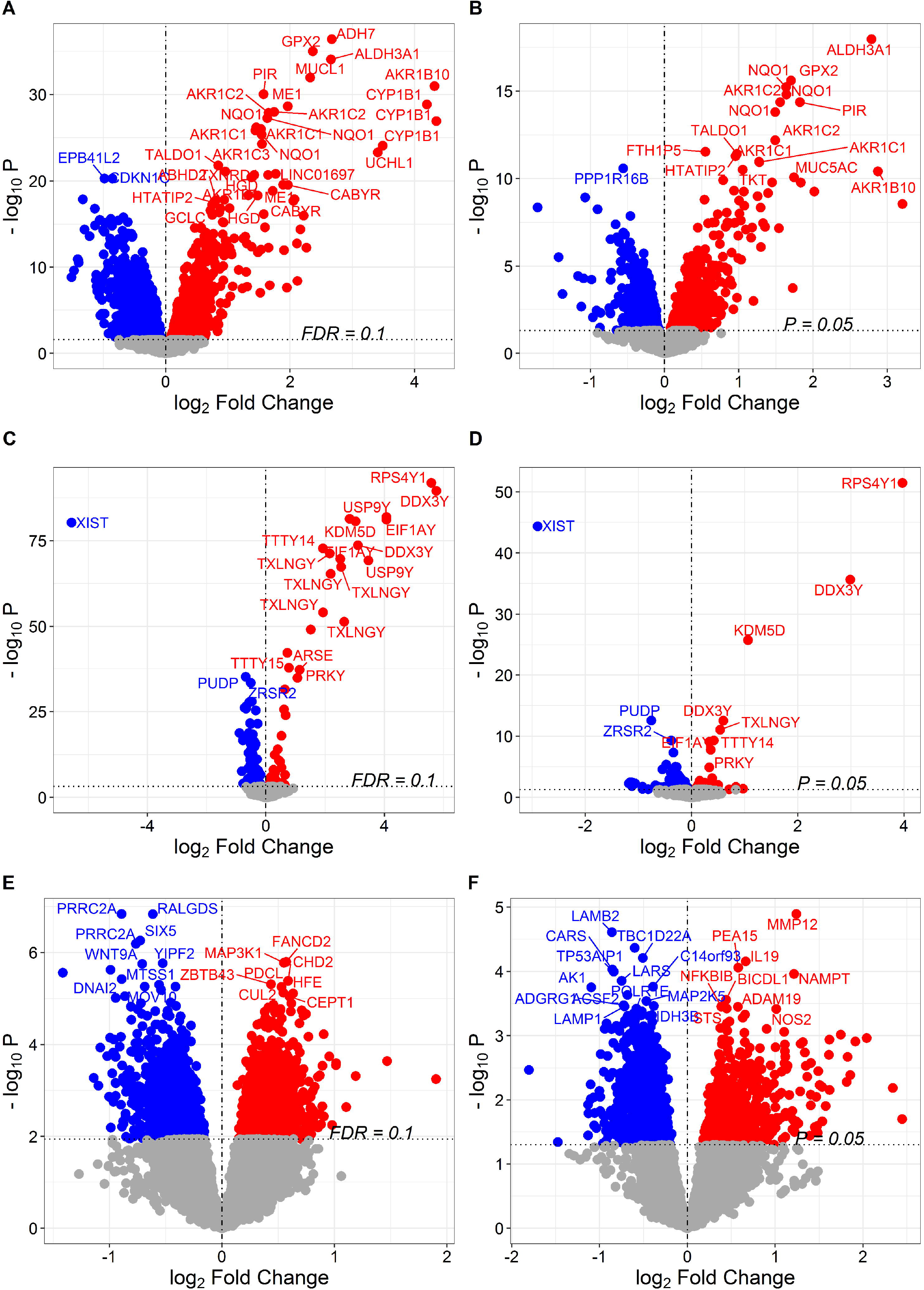
Volcano plots showing probe-phenotype associations. **(A)** Probe associations for smoking status in the discovery set. **(B)** Probe associations for smoking status in the replication set. **(C)** Probe associations for sex in the discovery set. **(D)** Probe associations for sex in the replication set. **(E)** Probe associations for sex-by-smoking interaction in the discovery set. **(F)** Probe associations for sex-by-smoking interaction in the discovery set. Probes shown in red were up-regulated in current smokers and probes shown in blue were down-regulated in current smokers in **(A)** and **(B)**. Probes shown in red were up-regulated in males and probes shown in blue were down-regulated in males in **(C)** and **(D)**. Probes shown in red exhibit a positive sex-by-smoking interaction in male current smokers while the probes shown in blue exhibit a negative sex-by-smoking interaction in male current smokers in **(E)** and **(F)**.

A total of 169 probes, corresponding to 100 genes were differentially expressed between males and females after adjustment for smoking status, age, ethnicity and pack-years (FDR < 0.1) and 27 of these genes were replicated in the replication set -(P < 0.05) (**Figure 3C and 3D**). Four of the replicated genes, *CCND2, FABP5, NLRP2*, and *SERPINB3*, are on the autosome. *CCND2* and *NLRP2* were down-regulated in males while *FABP5* and *SERPINB3* were up-regulated in males. The top three genes differentially expressed between males and females werewere *RPS4Y1* (FDR=3.98E-88), *DDX3Y* (FDR=4.33E-86) and *EIF1AY* (FDR=6.79E-82), which arelocated on the Y-chromosome (all three genes were up-regulated in males). Additional evaluation using GSE37147 replicated 22 out of the 27 replicated genes, including *CCND2* and *NLRP2*.

**Figure 3.**
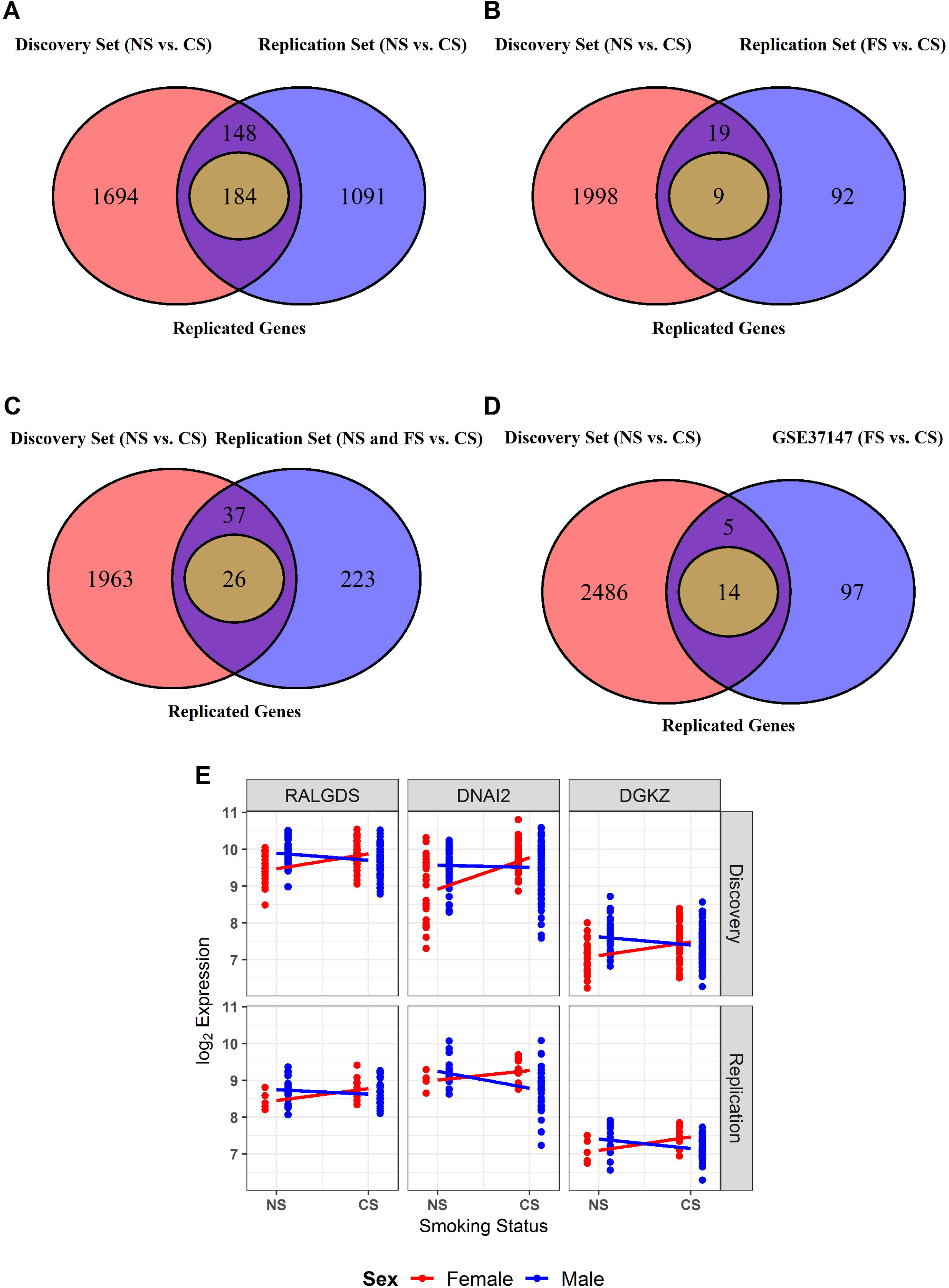
Replication of genes that were associated with sex-by-smoking interaction. **(A)** Venn diagram showing the number of associated genes in the discovery set (NS vs. CS) and in the replication set (NS vs. CS). **(B)** Venn diagram showing the number of associated genes in the discovery set (NS vs. CS) and in the replication set (NS and FS combined together vs. CS). **(C)** Venn diagram showing the number of associated genes in the discovery set (NS vs. CS) and in the replication set (FS vs. CS). **(D)** Venn diagram showing the number of associated genes in the discovery set (NS vs. CS) and in GSE37147 (FS vs. CS). **(E)** Gene expression of the top 3 replicated genes for sex-by-smoking interaction in the discovery and the replication set. Abbreviations used in the figure: NS = Never Smoker, FS = Former Smoker, CS = Current Smoker.

The complete lists of genes that were significantly associated with smoking status and sex are shown in **Supplementary Table 2** and **Supplementary Table 3**, respectively.

### Sexually dimorphic gene expression changes in response to smoking

Having identified genes differentially expressed according to sex and smoking status, we next identified sexually dimorphic genes in response to smoking. In the discovery set, 3,416 probes, corresponding to 2,654 genes were associated with sex-by-smoking interaction -(FDR < 0.1) (**Figure 2E**). In the replication set, 1,737 probes, corresponding to 1,423 genes, were associated with sex-by-smoking interaction - (P < 0.05) (**Figure 2F**). A total of 184 genes showed differential expression between never and current smokers in both datasets and within the same direction (**Figure 3A**). The top three replicated genes were *RALGDS* (FDR=2.19E-03), *DNAI2* (FDR=9.84E-03) and *DGKZ* (FDR=1.48E-03). The sexually dimorphic genes in response to smoking were not restricted to the XY chromosome; 96.5% of the associated genes in the discovery set and 94.0% of the 184 replicated genes were located on the autosome (**Figures 4A and 4B**). The lists of probes that were associated with the sex-by-smoking interaction term are shown in **Supplementary Table 4** in the discovery and the replication sets. Interestingly, the number of replicated genes decreased to 26 when we combined the never and former smokers together and compared them with the current smokers in the replication set, despite the increase in sample size (**Figure 3B**). The number of replicated genes decreased to 9 if we compared the former smokers to the current smokers in the replication set (**Figure 3C**). Similarly, when GSE37147 (containing current and former smokers but no never-smokers) was used as the replication set, only 14 genes were replicated (**Figure 3D**). **Figure 3E** shows the expression of the top three replicated genes in both the discovery and the replication set (never vs. current smokers). If we use GSE37147 to further replicate the 184 replicated genes, only one gene, *PLCB2*, was also replicated in GSE37147.

**Figure 4.**
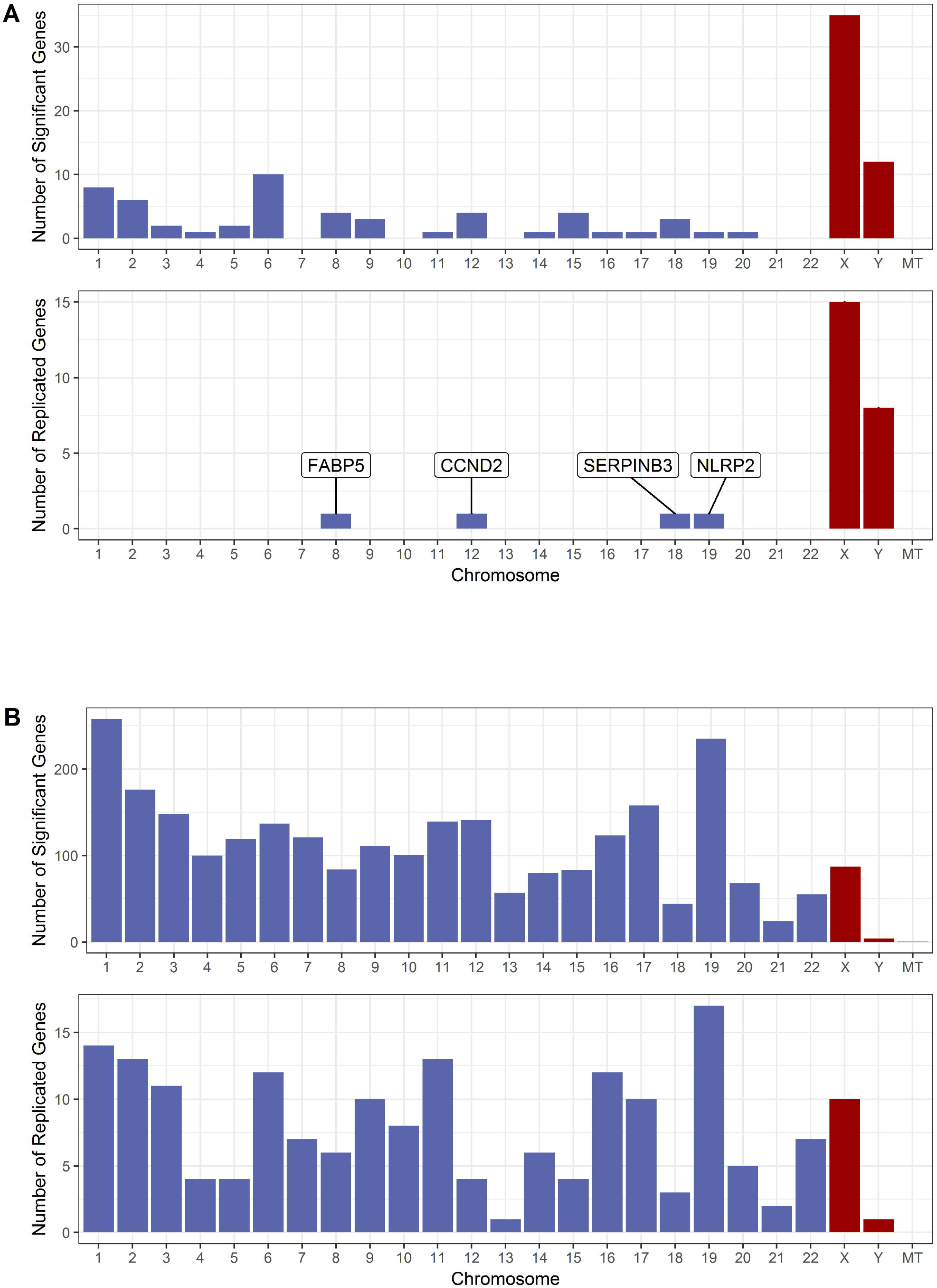
Chromosome distribution of genes that were associated with sex or sex-by-smoking interaction. **(A)** Top: Chromosome distribution of genes associated with sex in the discovery set (FDR < 0.1). Bottom: Chromosome distribution of the replicated genes for sex. **(B)** Top: Chromosome distribution of genes associated with sex-by-smoking interaction in the discovery set (FDR < 0.1). Bottom: Chromosome distribution of the replicated genes for sex-by-smoking interaction. Abbreviation used in the figure: MT=mitochondrion.

### Module-level associations

Based on the expression of 12,537 genes in the discovery set, 37 gene modules were constructed. The module sizes ranged from 17 genes in the “yellowgreen” module to 3,822 genes in the “turquoise” module. Eleven modules were significantly associated with smoking status (never vs. current smokers) after adjusting for age, sex, ethnicity and pack-years of smoking (FDR < 0.1) in the discovery set. Four of these modules, “darkgreen”, “greenyellow”, “grey60”, and “midnightblue”, were successfully replicated in the replication set (never and former vs. current smokers) at a threshold of P < 0.05.

Only the “yellowgreen” module was associated with sex after -(FDR < 0.1) in the discovery set and this module was also replicated in the replication set at a threshold of P < 0.05.

Thirteen modules were associated with sex-by-smoking interaction -in the discovery set (FDR < 0.1). Three modules, “blue”, “darkmagenta”, and “yellowgreen”, were successfully replicated in the replication set (P < 0.05). The “darkmagenta” and the “yellowgreen” modules were relatively small in size with 18 and 17 genes while the “blue” module included 1860 genes. The module-phenotype association along with the module sizes for the replicated modules are shown in **Figure 5**. The associations between the modules and the phenotypes are shown in **Supplementary Table 5**.

**Figure 5.**
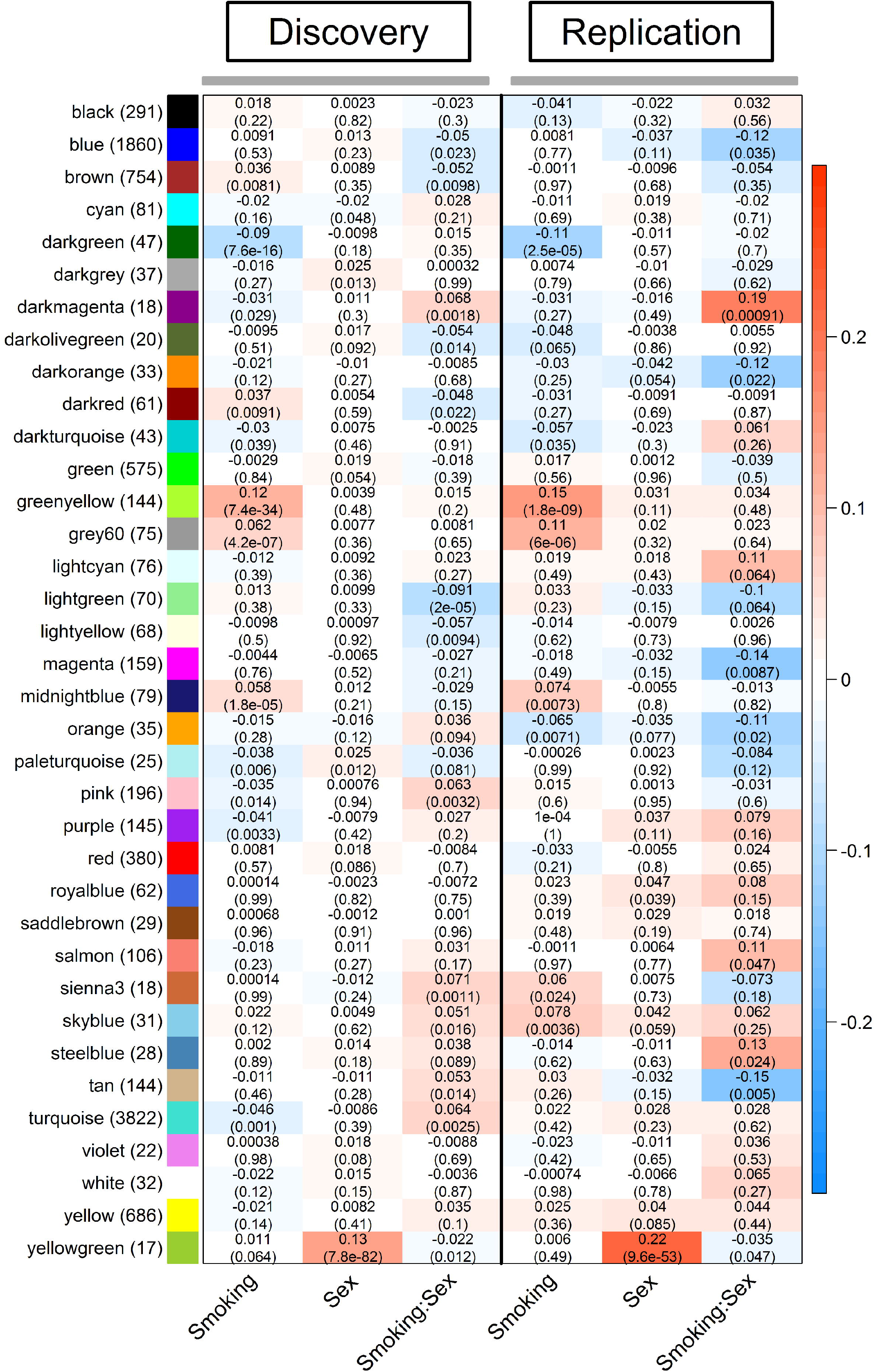
Module-level associations in the discovery and replication sets. The rows represent the gene modules and the sizes of the modules are shown within the brackets next to the module names. The columns represent the phenotypes. In each cell, the number at the top is the linear regression coefficient and the number in the brackets is the corresponding p-value. The color of the cell is proportional to the regression coefficient. From column 1 to column 3 of both the discovery and the replication sets, the red color indicates 1) up-regulation in current smokers, 2) up-regulation in male subjects, and 3) a positive sex-by-smoking interaction effect in male current smokers.

### Enrichment in Biological Processes

An FDR cutoff of 0.1 was used to identify Gene Ontology (GO) biological processes that were associated with the replicated modules. Among modules that were associated with smoking status, the “darkgreen” module genes were associated with protein complex assembly and the top genes with the highest MM in this module were *NPAS3, CYP4X1* and *PGRMC1*. The “greenyellow” module genes were associated with metabolic processes, responses to oxidative stress and homeostasis and the top members of this module were *AKR1B10, GPX2, AKR1C2*. The “midnightblue” module gene were related to cornification, protein glycosylation and viral entry into host cell. The top members of the “midnightblue” module are *ST14, HM13, TMPRSS4*. No significant pathways were enriched in the “grey60” module and the top members of this module were *CEACAM5, GALE* and *AGR2*.

Of the three modules associated with sex-by-smoking interaction, all the genes in the “yellowgreen” module belonged to the X and Y chromosomes and showed enrichment in translational initiation, protein targeting to endoplasmic reticulum and histone demethylation. The top members of the “yellowgreen” module were *DDX3Y, KDM5D* and *USP9Y*. This module was also associated with sex. For the remaining two modules, the “darkmagenta” module showed strong association with defense responses to virus, regulation of innate immune response, membrane fusion and Th2 cell cytokine production. The genes with the highest MM in the “darkmagenta module” are *DDX60, SAMD9* and *IFI44*. The “blue” module was related to autophagy, biological processes of mitochondrion, cellular response to oxidative stress and negative regulation of the MAPK signaling cascade. The top members of the “blue” module are *RALY, UQCRC1* and *MAP2K2*. **Table 2** shows the top GO biological processes enriched in the replicated modules and **Supplementary Table 6** shows the complete enrichment results for the replicated modules. **Supplementary Table 7** shows the list of top 10 genes with the highest membership in each module.

**Table 2.**
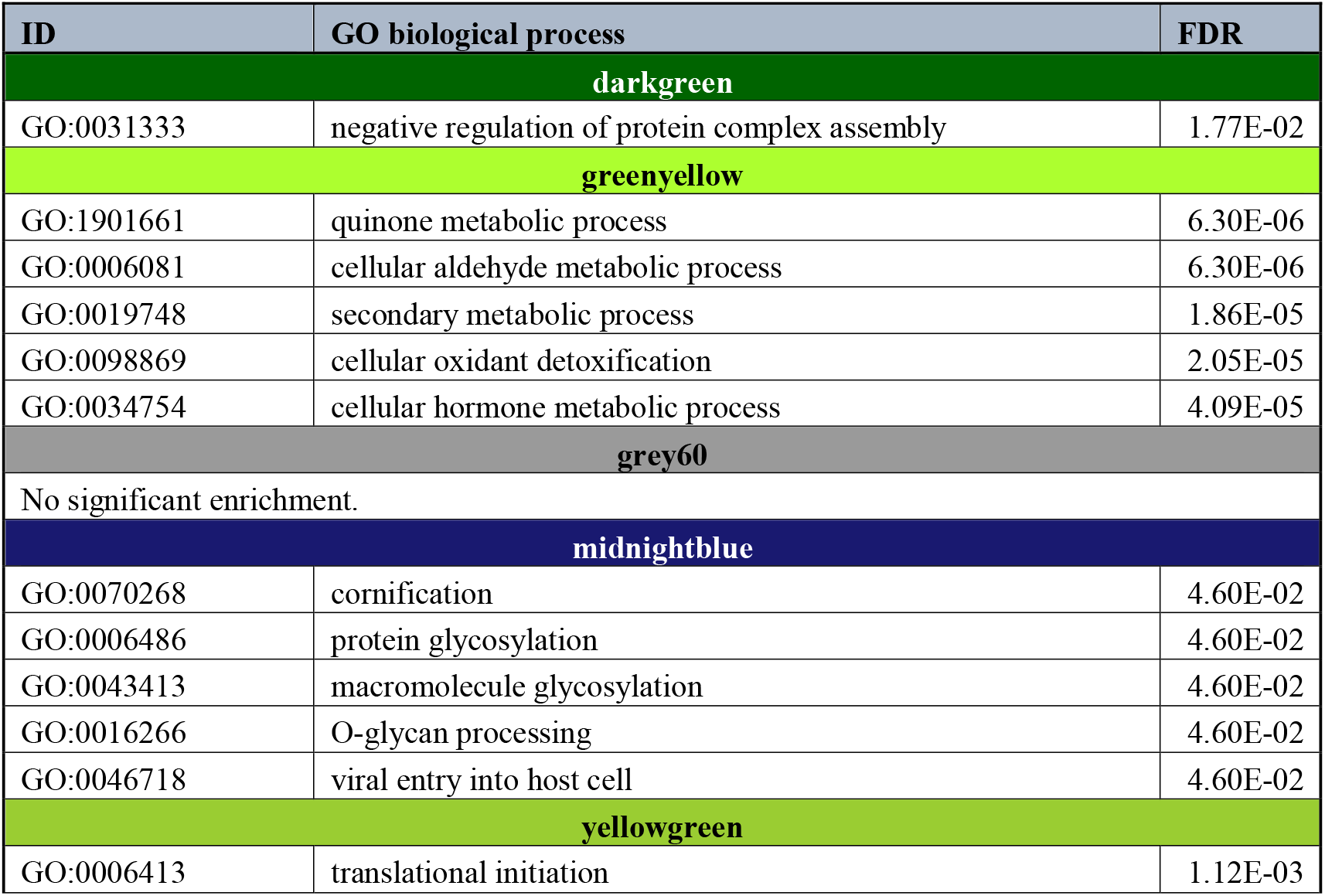
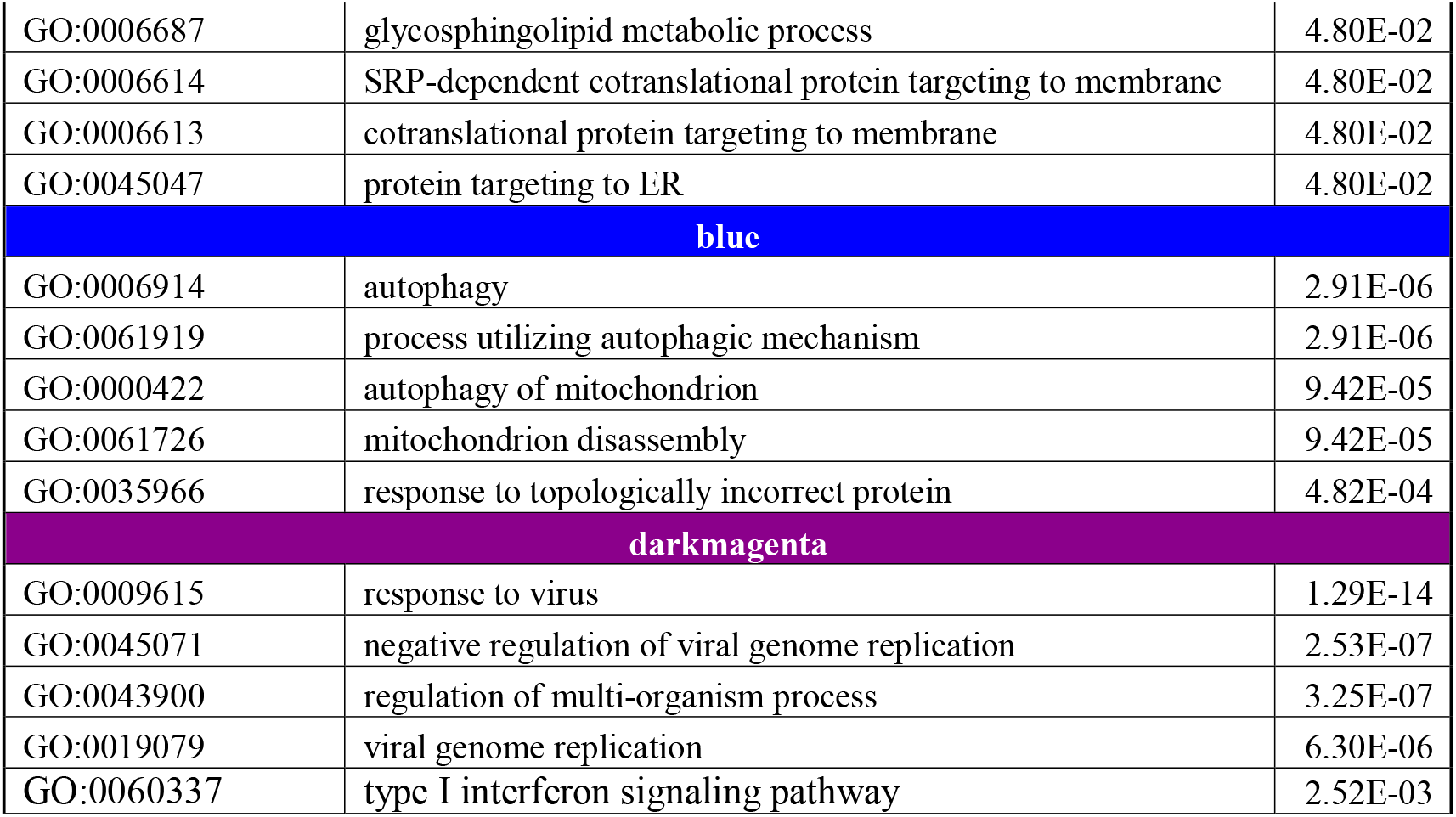
Top GO biological processes enriched in each replicated module.

| ID | GO biological process | FDR |
| --- | --- | --- |
| **darkgreen** |  |  |
| GO:0031333 | negative regulation of protein complex assembly | 1.77E-02 |
| **greenyellow** |  |  |
| GO:1901661 | quinone metabolic process | 6.30E-06 |
| GO:0006081 | cellular aldehyde metabolic process | 6.30E-06 |
| GO:0019748 | secondary metabolic process | 1.86E-05 |
| GO:0098869 | cellular oxidant detoxification | 2.05E-05 |
| GO:0034754 | cellular hormone metabolic process | 4.09E-05 |
| **grey60** |  |  |
| No significant enrichment. |  |  |
| **midnightblue** |  |  |
| GO:0070268 | cornification | 4.60E-02 |
| GO:0006486 | protein glycosylation | 4.60E-02 |
| GO:0043413 | macromolecule glycosylation | 4.60E-02 |
| GO:0016266 | O-glycan processing | 4.60E-02 |
| GO:0046718 | viral entry into host cell | 4.60E-02 |
| **yellowgreen** |  |  |
| GO:0006413 | translational initiation | 1.12E-03 |
| GO:0006687 | glycosphingolipid metabolic process | 4.80E-02 |
| GO:0006614 | SRP-dependent cotranslational protein targeting to membrane | 4.80E-02 |
| GO:0006613 | cotranslational protein targeting to membrane | 4.80E-02 |
| GO:0045047 | protein targeting to ER | 4.80E-02 |
| **blue** |  |  |
| GO:0006914 | autophagy | 2.91E-06 |
| GO:0061919 | process utilizing autophagic mechanism | 2.91E-06 |
| GO:0000422 | autophagy of mitochondrion | 9.42E-05 |
| GO:0061726 | mitochondrion disassembly | 9.42E-05 |
| GO:0035966 | response to topologically incorrect protein | 4.82E-04 |
| **darkmagenta** |  |  |
| GO:0009615 | response to virus | 1.29E-14 |
| GO:0045071 | negative regulation of viral genome replication | 2.53E-07 |
| GO:0043900 | regulation of multi-organism process | 3.25E-07 |
| GO:0019079 | viral genome replication | 6.30E-06 |
| GO:0060337 | type I interferon signaling pathway | 2.52E-03 |

In **Figure 6**, the gene expression changes for genes in the significant pathways: autophagy and response to virus are shown in males and females separately in response to smoking. While genes in autophagy pathway show little change in males, they were significantly up-regulated in females in response to smoking. On the other hand, genes in the viral response pathway showed little change in males but were significantly down-regulated in females in response to smoking.

**Figure 6.**
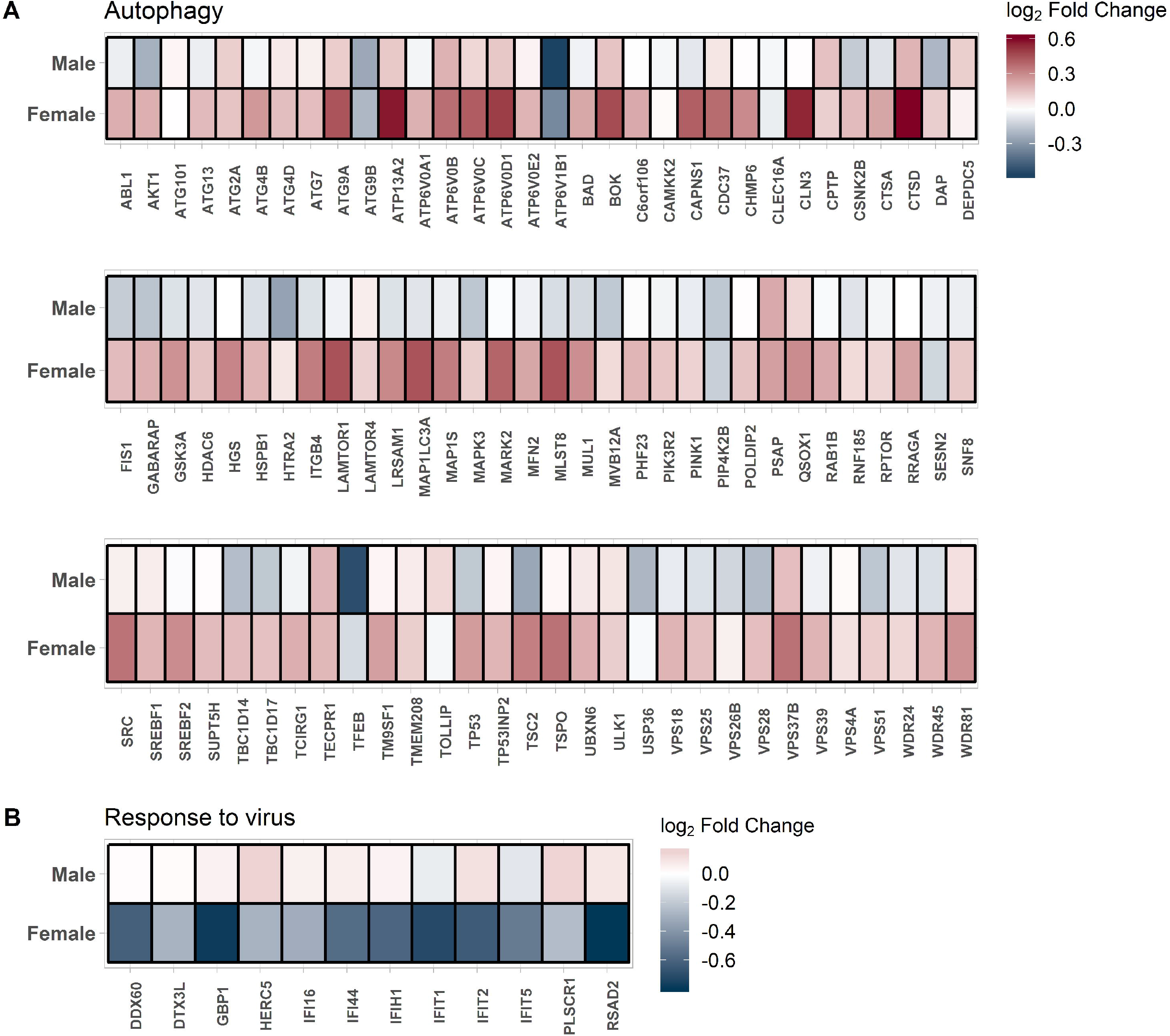
Sexually dimorphic effect of smoking on gene expression of genes in the top GO biological process in **(A)** the “blue” module and **(B)** in the “darkmagenta” module. The red color indicates up-regulation in current smokers and the blue color indicates down-regulation in current smokers.

### Enrichment in transcription factor binding site (TFBS)

An FDR cutoff of 0.05 and an odds ratio > 2 were used to identify TFBSs that were enriched in the replicated modules. For modules that were associated with smoking status, 13 TFBSs were enriched in the “darkgreen” module, 8 TFBSs were enriched in the “greenyellow” module, no TFBSs were enriched in the “grey60” module and 1 TFBS was enriched in the “midnightblue” module. No TFBS was enriched in the “yellowgreen” module which was associated with sex and sex-by-smoking interaction. For the remaining two modules that were associated with sex-by-smoking interaction, 72 TFBSs were enriched in the “blue” module including estrogen-related receptor alpha (*ESRRA*), and 7 TFBSs were enriched in the “darkmagenta” module. **Supplementary Table 8** shows the complete list of enriched TFBSs in each module.

## DISCUSSION

Studying the transcriptome is a very powerful hypothesis-free approach to elucidate the molecular mechanisms underlying diseases and phenotypes. In this study, we analyzed the differential transcriptomics of human airway epithelium at baseline and in response to smoking between males and females u. and found that 1) smoking has a large and reproducible effect on gene expression profiles; 2) the smoking effects on the transcriptome of human airways are modified by the underlying sex of the individual; 3) the sexually dimorphic smoking genes are common throughout the autosome and not restricted to sex chromosomes; and 4) the sexually dimorphic smoking genes are enriched in biological processes that are related to virus, anti-viral responses, type 1 interferon signaling, and autophagy.

There is increasing interest in understanding sexual dimorphisms across organs and tissues. In the study of Melé *et al.*^17^, the authors investigated sexual dimorphism in gene expression data of 1,641 samples from 175 individuals representing 43 sites and identified 753 genes that had tissue specific sex-biased expression (FDR < 0.05). Chen *et al.*^18^ used a much larger sample size (compared to Melé *et al.*) and investigated sexual dimorphism in gene expression data from the GTEx dataset: they evaluated 8,716 human transcriptomes collected from 31 tissues in 549 individuals of whom 188 were females (34%). The authors observed sexually dimorphic gene expression for 10% of the genes across autosomes in most tissues including the lung. The most sexually dimorphic gene expression were noted in the breast, skin, thyroid, brain, and adipose tissues while the least dimorphic was the gastrointestinal tract^18^. Multiple other studies in single cell or tissue types have confirmed to varying extents the large transcription differences between the two sexes^19–22^. Similar patterns emerged from these studies including the presence of large and perhaps underestimated differences in the transcriptome between males and females present throughout the autosomes and not restricted to sex chromosomes. Extrapolating findings from the published studies (even the large ones) to respiratory diseases is challenging since they do not contain data on airway epithelium and lack essential phenotypic information such as smoking. To our knowledge, the current study is the first to report sexually dimorphic gene expression changes in human airway epithelium in response to cigarette smoking. The airway epithelium is the initial site of injury in airway diseases such as COPD and is an important mediator of the host’s immune response to inhaled toxins, allergens and medications ^23^.

We found that smoking differentially altered two main pathways in men versus women. These included those related to host’s response to viruses (e.g. type 1 interferon signaling) and autophagy. In females, our study revealed that most autophagy genes were up-regulated in response to smoking compared to males where these genes were either down-regulated or only slightly up-regulated (**Figure 6A**). Autophagy is a physiological process that maintains homeostatic removal, degradation, and recycling of damaged proteins in response to physiological (e.g. development and immunity) and even pathological stressors such as oxidative stress ^24^. Increased autophagy has been reported in the lungs of COPD patients, in cigarette smoke-exposed mice as well as in airway epithelial cells exposed to cigarette smoke extracts in vitro ^25,26^. The disruption of autophagy caused by smoking may lead to epithelial loss of function and cell death^27^ which may adversely impact physiologic processes such as mucociliary clearance, and cellular senescence, which together would increase the risk of disease ^24,28,29^. Interestingly, sexual dimorphism in autophagy has been demonstrated in the spinal cord and skeletal muscles in mice ^30^ as well as the human hippocampus ^31^.

The other sexually dimorphic pathway in response in smoking was type 1 interferon signaling and response to virus. Genes in this pathway were down-regulated in females in response to smoking while being neutral or slightly up-regulated in males (**Figure 6B**). Previous studies have demonstrated that smoking affects the lung epithelium’s ability to produce the early inductive, and amplification phases of the type I interferon response and decreases the ability to elicit an antiviral response ^32,33^. Furthermore, these effects were reversible and are almost completely abrogated with the antioxidant glutathione^32^. The sexual dimorphism in immune response to viruses is well described (reviewed in ^34,35^). The reasons underlying this sexually dimorphic response is attributed to sex hormones, sex chromosomes linked genes involved in viral response or genes on autosomes that seem to be regulated by sex chromosome linked genes ^34^. Certain sexually dimorphic immune responses are however present throughout life, and not necessarily between the years of puberty and reproductive senescence, suggesting that both genes and hormones are involved ^36^. In general, females are thought to mount a stronger immune response to viral infections compared to males due to a more robust humoral and cellular immune response resulting in higher male mortality observed in epidemiological studies^37^. This is also consistent with observations that females have a higher risk to many autoimmune diseases ^38^. We observed an enrichment in many transcription factors for the autophagy module including the estrogen-related receptor alpha *ESRRA* (enrichment FDR = 7.02E-33) suggesting the biological differences are not only due to the hormonal effect. Further studies are needed to understand the clinical consequences of this sexually dimorphic virus response to smoking with regard to COPD risk, progression as well as exacerbations attributed to viral and bacterial infections. The finding of down-regulation of anti-viral response pathway genes in females in response to smoking is consistent with the epidemiological observations of higher exacerbation rate in females compared to males^8^.

We have used both gene and network level analyses in this study. The network level approach to transcriptomic data is biologically intuitive since it takes into account the phenomenon of gene co-expression. We have previously used network approach using WGCNA to identify a reproducible signature for lung function in peripheral blood^39^. The approach allows the identification of hub genes that reflect the biological processes of their corresponding modules. For example, the “hub” gene for the virus response enriched module “darkmagenta” is *DDX60* (DExD/H-box helicase 60) gene; a gene that is involved in antiviral response ^40^.

The definition of smoking has a big effect on discovery and replication of sexually dimorphic gene expression in response to smoking. When we included the data from GSE37147 (Sterling et al.) which included former and current smokers (but no never smokers) for replication, although the sample size is large (n=238), the replication was poor and hence was not used for replication (**Figure 3D**). Similarly, when we included both former and never smokers in our replication cohort vs. current smokers, and despite the increase in sample size, fewer genes were actually replicated compared to when we only included never vs. current smokers (**Figure 3A-3C**). This is consistent with previous findings from our group highlighting the importance of case definition of smokers on gene signature discovery^41^.

This study has a number of limitations. First, the sample size, although large, may not be sufficient to detect and replicate genes with smaller effects. Second, we used self-reported smoking status based on available data on GEO but self-reports may not accurately reflect smoking habits^41^. Third, we studied changes in airway epithelium given its role in the pathogenesis of lung diseases, but smoking may also affect other cell types such as immune and infiltrating cells in the airways. Finally, changes at gene expression may not necessarily reflect changes in protein levels or activities.

In summary, our study demonstrated that while smoking has a large effect on the transcriptome of airway epithelium, it differentially affects the viral response and autophagy pathways in female smokers compared to males. Such differential expression may explain the increased risk for disease and exacerbation among female smokers. These findings will stimulate mechanistic studies on how sex-specific genes impact risk to smoking related diseases such as COPD.

## METHODS

### Data collection

The search terms “airway epithelium” was used to identify gene expression studies of human airway epithelium gene expression from the Gene Expression Omnibus (GEO). After excluding irrelevant studies such as cancer, virus infection, influenza and injury, we have identified 18 studies with phenotypic information on subjects’ sex, smoking status and pack-years of smoking. Information on these 18 studies is shown in **Supplementary Table 1**. Of the studies identified, 16 were reported by the same lab and generated using the same microarray platform (U133A Plus 2.0 array). These datasets were combined together to form the “discovery” set. Phenotypic information such as sex, smoking status, age, COPD status, ethnicity and pack-years were available for 211 subjects (never smokers n=68; current smokers n=143) after removing duplicate data and outliers. GSE7895, which was generated using the U133A array, was used as the “replication” set consisting of 21 never smokers, 52 current smokers, and 31 former smokers. Smoking status, age and pack years were available from GEO for GSE7895 but the sex information was missing which was imputed for this cohort based on gene expression data from the discovery cohort (see below). GSE37147, which was generated using the GeneChip Human Gene 1.0 ST Array, was used as an additional replication cohort. Smoking status, sex, pack-years and lung function measurements of forced expiratory volume in 1 second (FEV_1_) and FEV_1_/forced vital capacity (FEV_1_/FVC) were available for 99 current smokers and 139 former smokers. **Figure 1** shows a flowchart of the overall study design.

### Data processing

Raw data (.CEL files) were downloaded from GEO for the discovery set, the replication set and GSE37147. Since the 16 studies of the discovery set contained overlapping samples, we carefully examined the phenotypic information of each sample to remove biological and technical replicates. No batch effect was detected among these studies using principal component analysis (**Supplementary Figure 1A**). The raw data of the discovery set were then normalized together using the Robust Multi-array Average (RMA) method. Similar to the discovery set, the replication set was also normalized using the RMA method (**Supplementary Figure 1B**). The presence of outliers was assessed using a principal component analysis before and after normalization for both the discovery and replication sets. Three outliers were observed and removed from the discovery set (**Supplementary Figure 1C**) while no obvious outliers were observed in the replication set (**Supplementary Figure 1D**). GSE37147 was also processed similarly and no obvious batch effect or outliers were detected. Large differences in the gene expression profiles were observed among the discovery set, the replication set and GSE37147 (**Supplementary Figure 1E**). All data processing and statistical analysis were performed within the R statistical computing environment (version 3.5.0).

### Prediction of sex in the replication set

Since the sex information of the replication set was not available from GEO, thus we imputed the sex of subjects using gene expression of selected XY-chromosome probes. To identify the probes that best discriminated male from female subjects, elastic net-regularized logistic regression was used for probe selection. The discovery set was partitioned into a training dataset, which contained 70% of female subjects and an equal number of male subjects, and a test dataset, which contained the remaining participants. Models with parameters were built based on the training dataset. The area under the receiver operating characteristics curve (AUC) obtained from 100 repeats of a 10-fold cross-validation was used as the criterion to evaluate the performance of the models. The performance of the model with the highest AUC was validated in the testing dataset. Finally, the model was applied to the replication set to predict the sex information.

### Comparing the demographic characteristics

We performed Kruskal-Wallis H test followed by Dunn’s test with the Bonferroni correction to compare age and pack-years of smoking between males and females within different smoking status strata in the discovery set, the replication set and GSE37147.

### Modeling the effect of smoking and sex on gene expression

Linear regression was used to identify probes that were associated with smoking status, sex and sex-by-smoking interaction adjusting for potential covariates such as age, ethnicity and pack-years of smoking in both the discovery and replication sets. The analyses were performed using R package limma ^42^. The Benjamini-Hochberg method was used to control the false discovery rate (FDR) and to correct for multiple hypotheses testing.

### Weighted gene co-expression network analysis (WGCNA)

The R package WGCNA ^39^ was used to construct gene co-expression modules with genes that are highly correlated with each other. The workflow of WGCNA starts with creating a matrix of Pearson correlations between genes, and transforming these into an adjacency matrix through soft thresholding and then converting them into a topological overlap matrix (TOM)^43^. Modules are defined as groups of highly interconnected genes using the average linkage hierarchical clustering to group genes based on the topological overlap of their connectivity, followed by a dynamic cut-tree algorithm to cluster dendrogram branches into gene modules^44^. Each gene within a module will be assigned a Module Membership (MM) whose value ranges between 0 and 1 by relating the gene’s expression profile with the module eigengene determined by the first principal component of the gene expression profiles in that module. Genes with the highest MM are considered “hub” genes, which are thought to be central to the module.

Since the discovery set and the replication set were generated using different microarray platforms, we first summarized the probe-level data of both sets into gene-level data using R package Jetset ^45^ before performing WGCNA. Using the 12,537 genes of the discovery set, we identified 37 gene modules and each module was assigned a color. For each of the gene module, an eigengene, which is representative of the module expression profile, was obtained. Linear regression was used to identify module eigengenes that were associated with smoking status, sex, and sex-by-smoking interaction, and COPD status adjusting for ethnicity and pack-years of smoking. To replicate our findings for modules constructed in the discovery set, we calculated their eigengenes in the replication set and then tested for their relationships to smoking status, sex, and sex-by-smoking interaction using linear regression. Similar to the discovery set, the models were adjusted for potential covariates such as pack-years.

### Enrichment analysis

For gene modules that were successfully replicated in the replication set, we performed enrichment analyses using R package clusterProfiler ^46^ and TFEA.ChIP ^47^ to identify Gene Ontology (GO) biological processes and transcription factor binding sites that were enriched in the module genes.

## AUTHORS’ CONTRIBUTIONS

Conceived and designed the study: MO, TLH, JL, DDS

Gene expression data analyses: MO, CXY, HS, ID, CWTY, EKHK,

Wrote the manuscript: MO, DDS, JL

Discussed results and implications and commented on the manuscript at all stages: all co-authors.

All authors read and approved the final manuscript.

## ACKNOWLEDGEMENTS

M.O is a fellow of the Parker B. Francis Foundation and a Scholar of the Michael Smith Foundation for Health Research. D.D.S holds Canada Research Chair in COPD.

---

**Supplementary Figure 1.**
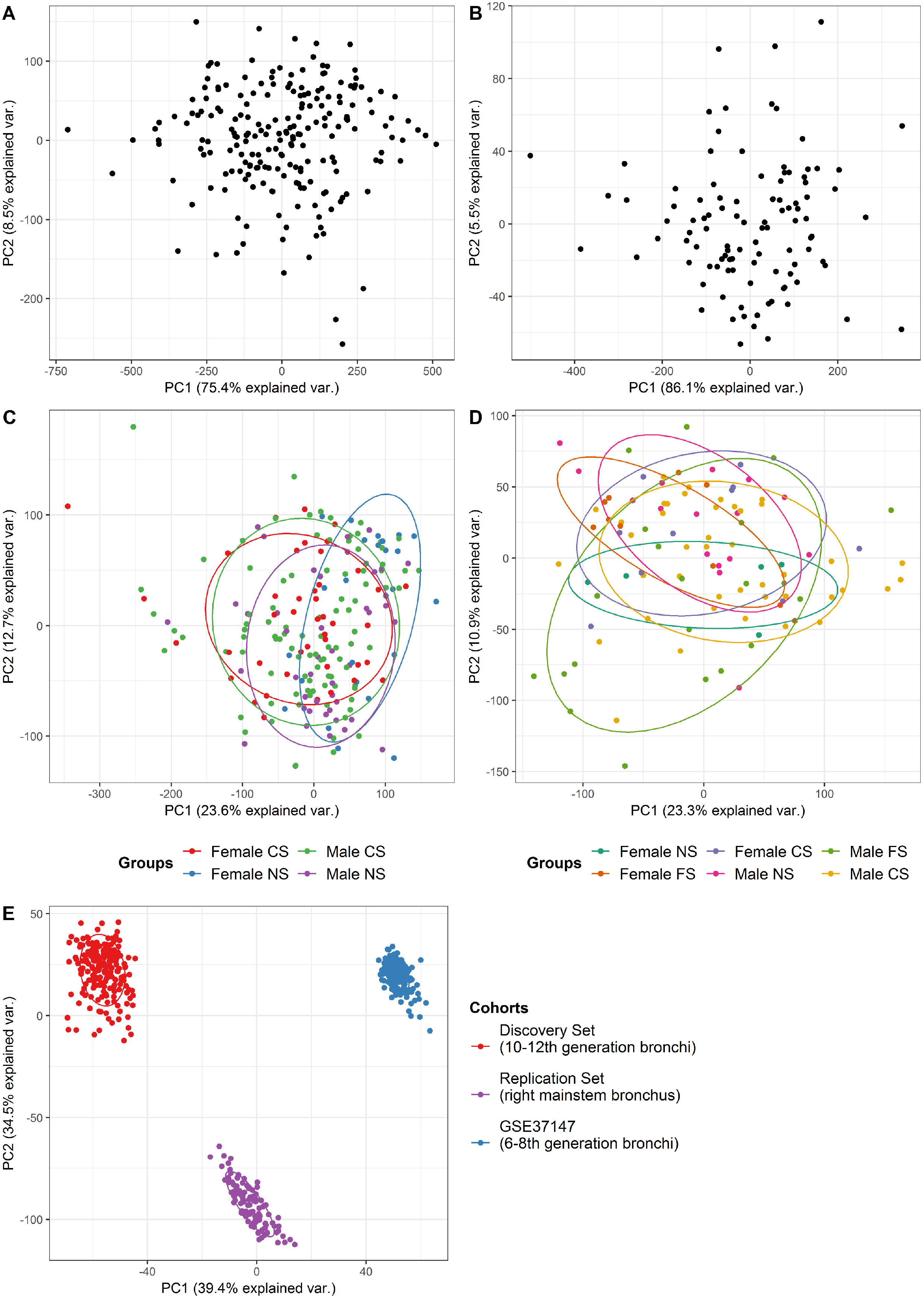
Assessing batch effect and outliers. **(A)** PCA plot on the raw data of the discovery set. **(B)** PCA plot on the raw data of the replication set. **(C)** PCA plot on the normalized data of the discovery set. **(D)** PCA plot on the normalized data of the replication set. **(E)** PCA plot on the combined data of the discovery set, the replication set and GSE37147. Abbreviations used in the figure: NS = Never Smoker, FS = Former Smoker, CS = Current Smoker.

**Supplementary Figure 2.**
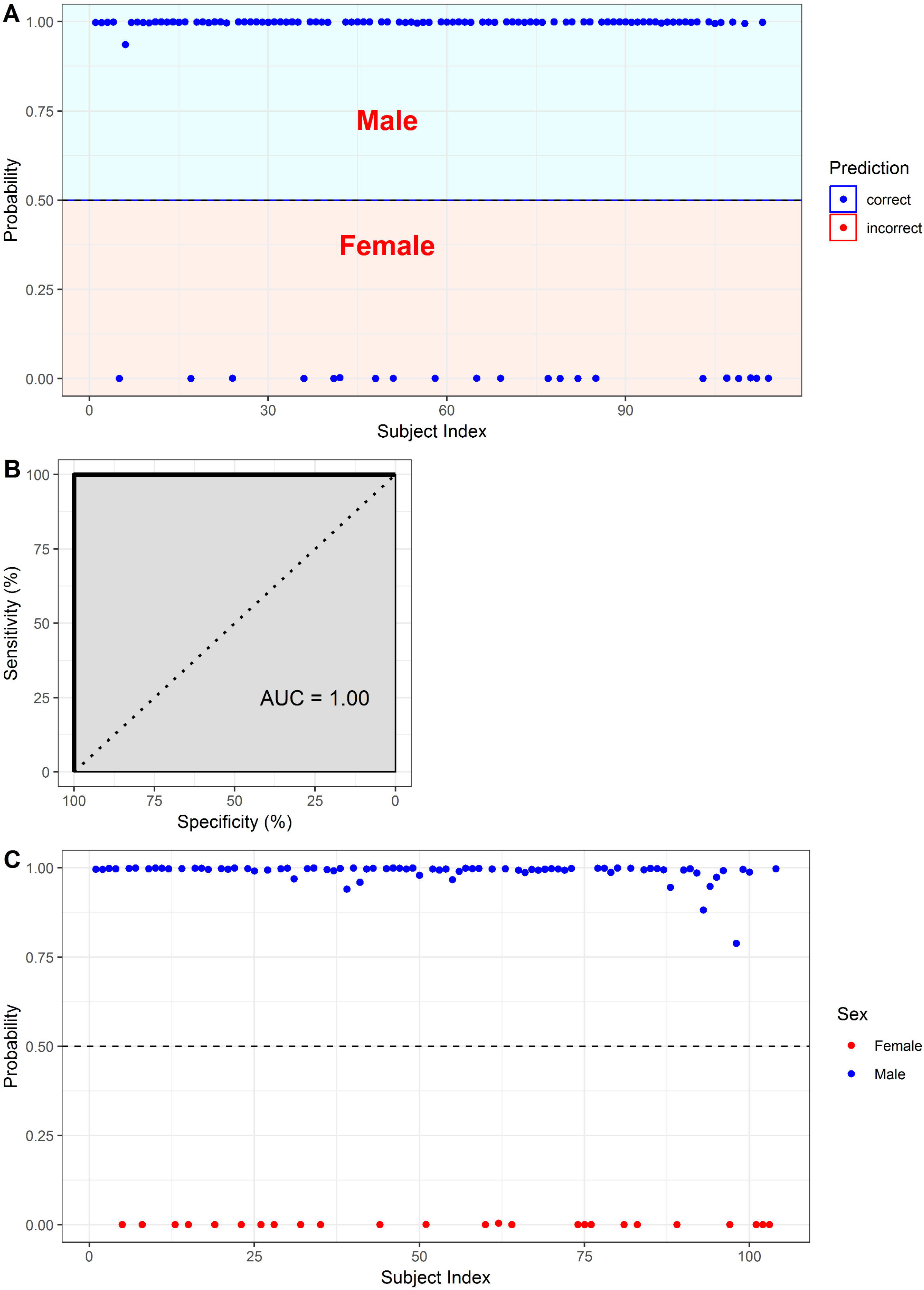
Prediction of sex. **(A)** Prediction of sex in the testing data of the discovery set. **(B)** Model performance. **(C)** Prediction of sex in the replication set.

## REFERENCES

1 Noguchi, E. I. et al. Evidence for Linkage Between Asthma/Atopy in Childhood and Chromosome 5q31-q33 in a Japanese Population. American Journal of Respiratory and Critical Care Medicine 156, 1390–1393, doi:10.1164/ajrccm.156.5.9702084 (1997).

2 Lozano, R. et al. Global and regional mortality from 235 causes of death for 20 age groups in 1990 and 2010: a systematic analysis for the Global Burden of Disease Study 2010. Lancet 380, 2095–2128, doi:10.1016/S0140-6736(12)61728-0 (2012).

3 Tager, I. B., Segal, M. R., Speizer, F. E. & Weiss, S. T. The natural history of forced expiratory volumes. Effect of cigarette smoking and respiratory symptoms. The American review of respiratory disease 138, 837–849, doi:10.1164/ajrccm/138.4.837 (1988).

4 Evans, J., Chen, Y., Camp, P. G., Bowie, D. M. & McRae, L. Estimating the prevalence of COPD in Canada: Reported diagnosis versus measured airflow obstruction. Health reports 25, 3–11 (2014).

5 Gan, W. Q., Man, S. F., Postma, D. S., Camp, P. & Sin, D. D. Female smokers beyond the perimenopausal period are at increased risk of chronic obstructive pulmonary disease: a systematic review and meta-analysis. Respiratory research 7, 52, doi:10.1186/1465-9921-7-52 (2006).

6 Amaral, A. F. S., Strachan, D. P., Burney, P. G. J. & Jarvis, D. L. Female Smokers Are at Greater Risk of Airflow Obstruction Than Male Smokers. UK Biobank. American journal of respiratory and critical care medicine 195, 1226–1235, doi:10.1164/rccm.201608-1545OC (2017).

7 Celli, B. et al. Sex differences in mortality and clinical expressions of patients with chronic obstructive pulmonary disease. The TORCH experience. American journal of respiratory and critical care medicine 183, 317–322, doi:10.1164/rccm.201004-0665OC (2011).

8 Celli, B. et al. Sex differences in mortality and clinical expressions of patients with chronic obstructive pulmonary disease. The TORCH experience. Am J Respir Crit Care Med 183, 317–322, doi:10.1164/rccm.201004-0665OC (2011).

9 Dransfield, M. T. et al. Gender differences in the severity of CT emphysema in COPD. Chest 132, 464–470, doi:10.1378/chest.07-0863 (2007).

10 Martinez, F. J. et al. Sex differences in severe pulmonary emphysema. American journal of respiratory and critical care medicine 176, 243–252, doi:10.1164/rccm.200606-828OC (2007).

11 Tam, A. et al. Sex Differences in Airway Remodeling in a Mouse Model of Chronic Obstructive Pulmonary Disease. Am J Respir Crit Care Med 193, 825–834, doi:10.1164/rccm.201503-0487OC (2016).

12 Benowitz, N. L., Lessov-Schlaggar, C. N., Swan, G. E. & Jacob, P., 3rd. Female sex and oral contraceptive use accelerate nicotine metabolism. Clinical pharmacology and therapeutics 79, 480–488, doi:10.1016/j.clpt.2006.01.008 (2006).

13 Sin, D. D., Cohen, S. B., Day, A., Coxson, H. & Pare, P. D. Understanding the biological differences in susceptibility to chronic obstructive pulmonary disease between men and women. Proceedings of the American Thoracic Society 4, 671–674, doi:10.1513/pats.200706-082SD (2007).

14 Van Winkle, L. S., Gunderson, A. D., Shimizu, J. A., Baker, G. L. & Brown, C. D. Gender differences in naphthalene metabolism and naphthalene-induced acute lung injury. American journal of physiology. Lung cellular and molecular physiology 282, L1122–1134, doi:10.1152/ajplung.00309.2001 (2002).

15 Tam, A. et al. The role of female hormones on lung function in chronic lung diseases. BMC women’s health 11, 24, doi:10.1186/1472-6874-11-24 (2011).

16 Tam, A., Wadsworth, S., Dorscheid, D., Man, S. F. & Sin, D. D. Estradiol increases mucus synthesis in bronchial epithelial cells. PloS one 9, e100633, doi:10.1371/journal.pone.0100633 (2014).

17 Mele, M. et al. Human genomics. The human transcriptome across tissues and individuals. Science 348, 660–665, doi:10.1126/science.aaa0355 (2015).

18 Chen, C.-Y. et al. Sexual dimorphism in gene expression and regulatory networks across human tissues. bioRxiv, doi:10.1101/082289 (2016).

19 Vawter, M. P. et al. Gender-Specific Gene Expression in Post-Mortem Human Brain: Localization to Sex Chromosomes. Neuropsychopharmacology: official publication of the American College of Neuropsychopharmacology 29, 373–384, doi:10.1038/sj.npp.1300337 (2004).

20 Cheng, C. & Kirkpatrick, M. Sex-Specific Selection and Sex-Biased Gene Expression in Humans and Flies. PLoS Genetics 12, e1006170, doi:10.1371/journal.pgen.1006170 (2016).

21 Trabzuni, D. et al. Widespread sex differences in gene expression and splicing in the adult human brain. 4, 2771, doi:10.1038/ncomms3771 https://www.nature.com/articles/ncomms3771#supplementary-information (2013).

22 Wcrling, D. M., Parikshak, N. N. & Geschwind, D. H. Gene expression in human brain implicates sexually dimorphic pathways in autism spectrum disorders. Nature Communications 7, 10717, doi:10.1038/ncomms10717 (2016).

23 Aghapour, M., Raee, P., Moghaddam, S. J., Hiemstra, P. S. & Heijink, I. H. Airway Epithelial Barrier Dysfunction in COPD: Role of Cigarette Smoke Exposure. American journal of respiratory cell and molecular biology, doi:10.1165/rcmb.2017-0200TR (2017).

24 Nyunoya, T. et al. Molecular processes that drive cigarette smoke-induced epithelial cell fate of the lung. American journal of respiratory cell and molecular biology 50, 471–482, doi:10.1165/rcmb.2013-0348TR (2014).

25 Chen, Z. H. et al. Egr-1 regulates autophagy in cigarette smoke-induced chronic obstructive pulmonary disease. PloS one 3, e3316, doi:10.1371/journal.pone.0003316 (2008).

26 Kim, H. P. et al. Autophagic proteins regulate cigarette smoke-induced apoptosis: protective role of heme oxygenase-1. Autophagy 4, 887–895 (2008).

27 Wang, G. et al. Role of OSGIN1 in mediating smoking-induced autophagy in the human airway epithelium. Autophagy 13, 1205–1220, doi:10.1080/15548627.2017.1301327 (2017).

28 Lam, H. C. et al. Histone deacetylase 6-mediated selective autophagy regulates COPD-associated cilia dysfunction. The Journal of clinical investigation 123, 5212–5230, doi:10.1172/jci69636 (2013).

29 Pallet, N. Cigarette smoke-induced autophagy: a deadly association? Journal of thoracic disease 9, 2228–2230, doi:10.21037/jtd.2017.06.132 (2017).

30 Oliv et al. Sex Differences in Constitutive Autophagy. BioMed Research International 2014, 5, doi: 10.1155/2014/652817 (2014).

31 Guebel, D. V. & Torres, N. V. Sexual Dimorphism and Aging in the Human Hyppocampus: Identification, Validation, and Impact of Differentially Expressed Genes by Factorial Microarray and Network Analysis. Frontiers in Aging Neuroscience 8, doi:10.3389/fnagi.2016.00229 (2016).

32 Bauer, C. M. T. et al. Cigarette Smoke Suppresses Type I Interferon-Mediated Antiviral Immunity in Lung Fibroblast and Epithelial Cells. Journal of Interferon & Cytokine Research 28, 167–179, doi:10.1089/jir.2007.0054 (2008).

33 Bauer, C. M. T., Morissette, M. C. & Stampfli, M. R. The influence of cigarette smoking on viral infections: translating bench science to impact COPD pathogenesis and acute exacerbations of COPD clinically. Chest 143, 196–206, doi:10.1378/chest.12-0930 (2013).

34 Ghosh, S. & Klein, R. S. Sex Drives Dimorphic Immune Responses to Viral Infections. J Immunol 198, 1782–1790, doi:10.4049/jimmunol.1601166 (2017).

35 Klein, S. L. & Flanagan, K. L. Sex differences in immune responses. Nature reviews. Immunology 16, 626–638, doi:10.1038/nri.2016.90 (2016).

36 Klein, S. L. & Flanagan, K. L. Sex differences in immune responses. Nature Reviews Immunology 16, 626, doi:10.1038/nri.2016.90 (2016).

37 Klein, S. L. Sex influences immune responses to viruses, and efficacy of prophylaxis and treatments for viral diseases. BioEssays : news and reviews in molecular, cellular and developmental biology 34, 1050–1059, doi:10.1002/bies.201200099 (2012).

38 McCombe, P. A., Greer, J. M. & Mackay, I. R. Sexual dimorphism in autoimmune disease. Curr Mol Med 9, 1058–1079 (2009).

39 Obeidat, M. e. et al. Network-based analysis reveals novel gene signatures in peripheral blood of patients with chronic obstructive pulmonary disease. Respiratory research 18, 72, doi: 10.1186/s12931-017-0558-1 (2017).

40 Oshiumi, H. et al. DDX60 Is Involved in RIG-I-Dependent and Independent Antiviral Responses, and Its Function Is Attenuated by Virus-Induced EGFR Activation. Cell Rep 11, 1193–1207, doi:10.1016/j.celrep.2015.04.047 (2015).

41 Obeidat, M. e. et al. The Effect of Different Case Definitions of Current Smoking on the Discovery of Smoking-Related Blood Gene Expression Signatures in Chronic Obstructive Pulmonary Disease. Nicotine & Tobacco Research 18, 1903–1909, doi:10.1093/ntr/ntw129 (2016).

42 Ritchie, M. E. et al. limma powers differential expression analyses for RNA-sequencing and microarray studies. Nucleic acids research 43, e47, doi:10.1093/nar/gkv007 (2015).

43 Yip, A. & Horvath, S. Gene network interconnectedness and the generalized topological overlap measure. BMC bioinformatics 8, 22 (2007).

44 Langfelder, P., Zhang, B. & Horvath, S. Defining clusters from a hierarchical cluster tree: the Dynamic Tree Cut package for R. Bioinformatics 24, 719–720, doi:10.1093/bioinformatics/btm563 (2008).

45 Li, Q., Birkbak, N. J., Gyorffy, B., Szallasi, Z. & Eklund, A. C. Jetset: selecting the optimal microarray probe set to represent a gene. BMC Bioinformatics 12, 474, doi: 10.1186/1471-210512-474 (2011).

46 Yu, G., Wang, L. G., Han, Y. & He, Q. Y. clusterProfiler: an R package for comparing biological themes among gene clusters. Omics 16, 284–287, doi:10.1089/omi.2011.0118 (2012).

47 Puente-Santamaria, L. & del Peso, L. TFEA.ChIP: A tool kit for transcription factor binding site enrichment analysis capitalizing on ChIP-seq datasets. bioRxiv, 303651, doi: 10.1101/303651 (2018).

